# Does participation in acoustic experiments improve welfare in captive animals? A case study of three grey seals (*Halichoerus grypus*)

**DOI:** 10.1101/2020.08.15.252460

**Authors:** Sara T. Ortiz, Alyssa Maxwell, Ariana Hernandez, Kirstin Anderson Hansen

**Affiliations:** Marine Biological Research Center, University of Southern Denmark, Hindsholmvej 11, 5300 Kerteminde, Denmark; Department of Biology, University of Southern Denmark, 5230 Odense, Denmark; Max Planck Institute for Ornithology, Eberhard-Gwinner-Strasse, 82319 Seewiesen, Germany; Institute for Terrestrial and Aquatic Wildlife Research, University of Veterinary Medicine, Foundation, Werftstr. 6, 25761 Büsum, Germany

**Keywords:** Welfare, Pinnipeds, Motivation, Personality, Enrichment

## Abstract

Both mental and physiological conditions determine the well-being state in an animal. Enrichment is a way to increase an animals’ well-being and may require problem solving through thinking, tolerance of ambiguity, openness, and intrinsic motivation. It is unclear if it is enriching when an animal participates in different types of research. Therefore, it is important to answer the question of whether research can be used as an enrichment tool in zoological facilities. Here, we examine if participation in psychophysical research affected the mental stimulation of three grey seals under human care. The effects varied amongst the three individuals that took part in the research, and indicated that their participation in the research task was dependent on their individual personalities and life history. Two seals indicated that their involvement in the research was positive and motivating, and therefore can be considered enriching. In comparison, the third seal displayed a tendency for frustration and low motivation. Our results indicate that research can be a powerful enrichment tool with animals that find research motivating.

## Introduction

A key topic over the last few years has been to ensure that the lives of captive animals are being properly enriched (Shepherdson, 1998). Positive experiences can be provided as a form of environmental enrichment (Boissy et al., 2007). Captive animals are mainly kept in settings that are less complex and smaller than in the wild (Buchanan-Smith, 1997; Chamove, 1989). Therefore, enrichment is necessary to guarantee that the animal receives the physical and mental stimulation that it would have in their natural habitats. Environmental enrichment has been shown to prominently reduce stress (Carlstead & Shepherdson, 2000), as well as raise the quality and diversify of the behavioral repertoires of captive animals (Makecha & Highfill, 2018). The goal of enrichment is to improve the target animal’s mental well-being and/or physical fitness, in other words, its welfare (Hunter et al., 2002; Swaisgood & Shepherdson, 2005). Previous studies have suggested that appropriate environmental enrichment enhances the physiological well-being of animals who have a positive human-animal relationship under human care (Vaicekauskaite et al., 2019). An animal’s life history traits, such as age, sex, and/or social status, influence the way they respond differently to environmental enrichment (Eskelinen et al., 2015). Therefore, it is crucial to examine individual responses in order to adapt the type of enrichment to the individual in question and not assume that individuals will respond the same to different forms of enrichment (Vaicekauskaite et al., 2019). If the enrichment is not adapted to the animal, both as a species or individual, it might not have the desired effect (Young, 2004).

Welfare of an individual could be defined as “The state of the animals’ body and mind as regards its attempts to cope with its environment”, referring to feelings and health as a part of it (Broom, 1986; Fraser, 2008). The word *personality* can also be used to describe the animal’s personal distinct and consistent behavioral reactions to different environmental variables, which remain stable over time and across situations (Hosey et al., 2013; Makecha & Highfill, 2018). Maintaining the animal’s welfare should be an essential part of keeping animals under human care (Makecha & Highfill, 2018). While there is a broad range of indicators of poor welfare, there are significantly less behaviors, and no physiological measures, used as indicators of good welfare (Bassett & Buchanan-Smith, 2007). Animal welfare has been shown to be positively affected by cognitive tasks (Clark, 2011, 2013; Moberg, 2000).

Scientific studies have a potential to improve an animal’s mental and physical stimulation, as well as increase our understanding of them (Clark, 2013; Hopper et al., 2016; Puppe et al., 2007; Yamanashi & Hayashi, 2011). Researchers can benefit from collaborating with both zoos and aquariums, as they reduce the cost of maintenance and housing of research animals, as well as provide a wide variety of species to investigate (Hopper, 2017). From this collaboration, zoos and aquariums may be able to provide their animals with new enrichment tools if participation in research is proven to be positively enriching. It is clear from previous studies that research participation in marine parks and zoos can give countless opportunities to provide meaningful enrichment to the participating animals.

Numerous studies have found positive effects of cognitive research on the welfare of the animals involved (Clark, 2011; Whitehouse et al., 2013; Yamanashi & Hayashi, 2011), but there are only a few studies about the positive effects of other types of research (Ruby & Buchanan-Smith, 2015). In this study we investigated if the participation of three captive grey seals (*Halichoerus grypus*) in a psychophysical research task can be considered enrichment over an extended period of time. Additionally, we examined if training history played a role in determining whether participating in research improved the welfare of the individuals.

## Materials and Methods

### Study subjects

The experimental animals (Table 1) were three captive-born male grey seals (*Halichoerus grypus*) housed at the Marine Biological Research Centre at the University of Southern Denmark (SDU) in Kerteminde, Denmark.

**Table 1.**
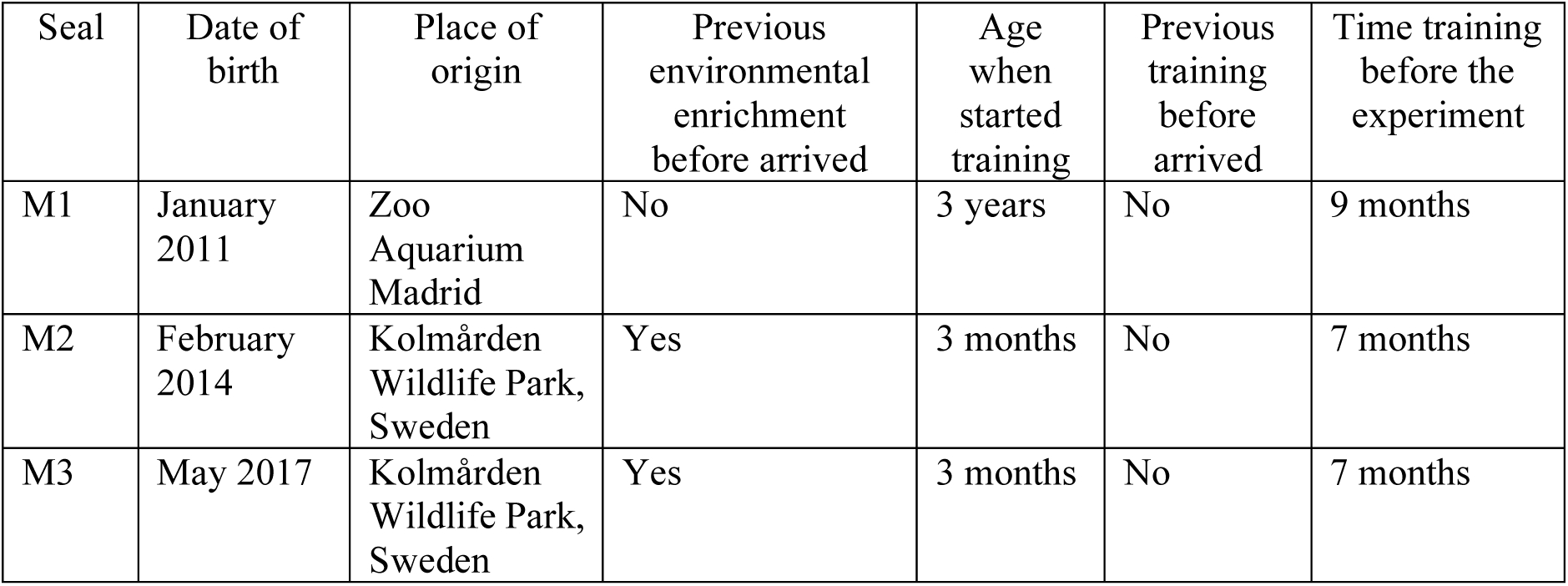
Enrichment and training history of the three individuals.

The seals were trained using operant conditioning with positive reinforcement 3-4 times a day. M1 began learning the research task when he was 3,5 years old. M2 and M3 began training for the research task when they were 4 months old.

### Housing

The animals were housed in an open-air enclosure located on the edge of a harbor. There were three circular housing pools: A: depth: 1.7 m, diameter: 4 m; B: depth: 1.7 m, diameter: 5 m; C: depth: 1.7 m, diameter 7 m (Figure 1). Pool C was an underground pool, which was better adapted for acoustic research. Each pool was surrounded by wood decking and metal mesh fencing and gates, which allowed the seals to have visual and acoustic contact with one another, as well as allowed the trainers to move and separate the subjects between pools.

**Figure 1.**
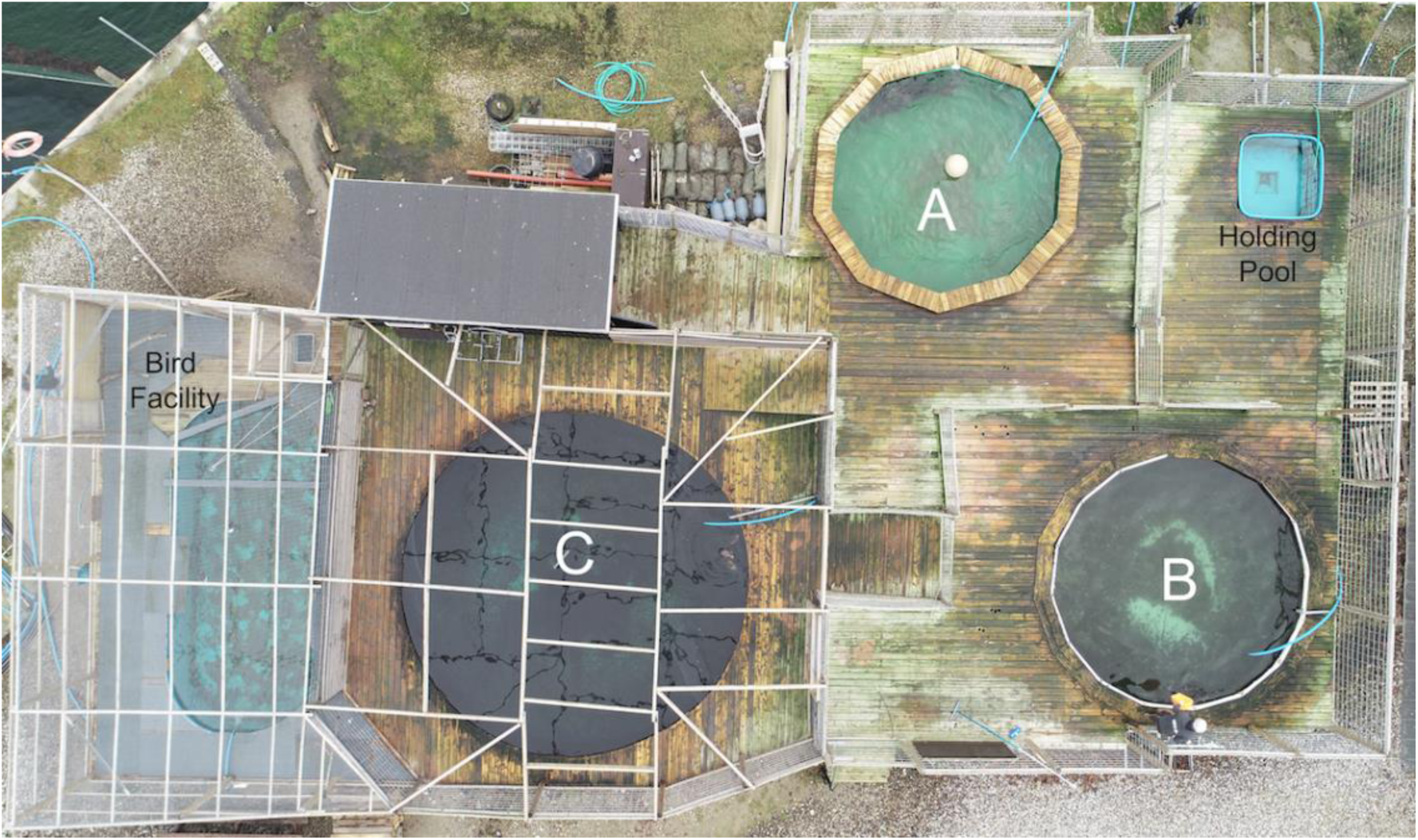
A schematic of the animal housing facility at the marine research center. Pools are labeled A-C (see methods).

### Husbandry

Separation was necessary during research and training sessions, as well as throughout the months of the breeding season due to the subjects being male, and the large size and weight difference between the three animals.

The seals’ diets were calculated to maintain a healthy weight, considering both the age and size of the animal, and the time of year. Research took place from January 2015 to June 2015 for M1, January 2015 to June 2018 for M2, and January 2018 to June 2018 for M3. The daily diet was divided amongst the 3-4 training sessions and used as reinforcement. On days when research was conducted, the diets were divided differently, so that 30-50% of their diet for the day was selected for the research session and the remaining 50-70% was divided among the remaining 2-3 training sessions. The diets consisted of a mixture of herring (*Clupea harengus*; 190 calories/100 grams), capelin (*Mallotus villosus*; 136 calories/100 grams) and sprat (*Sprattus sprattus*; 155 calories/100 grams). As daily enrichment, a variety of ‘toys’ were given to the animals every day. Toys ranged from surf boards, fire hoses, small to large buoys, boomer balls, and ice cubes with fish.

### Experimental procedure

The data used in this study was a part of a larger study investigating underwater hearing thresholds of the grey seal, thereby contributing to our understanding of the effects of anthropogenic noise on this species. In order to determine the animals’ hearing thresholds, the subjects were trained to participate in a psychophysical study with a GO/NOGO testing paradigm, using positive reinforcement procedures (Fay & Wilber, 1989; Gescheider, 2013; Green & Swets, 1966; Stebbins, 2013). Psychophysical testing procedures are commonly used to study how animals translate stimuli from its surrounding environment to sensations in their psychological domain (Gescheider, 2013), with the ultimate goal being to determine their sensory threshold (for example, see: Maxwell et al. (2017) and Hansen et al. (2017)). By using psychophysics, it is possible to determine the weakest detectable stimulus, known as a sensory threshold, as well as the minimum amount of stimulus required to have the animal produce a sensation (Gescheider, 2013). Two types of trials were used in this testing procedure: (1) signal present (GO) and (2) signal absent (NOGO). From these two trial types, there were four possible responses: 1) GO indicating a correct response to a signal present trial, 2) NOGO indicating a correct response to a signal absent trial, 3) MISS indicating an incorrect response to a signal present trial, and a FALSE ALARM indicating an incorrect response to a signal absent trial. In a session, the two different trial types were presented to the animal in a randomized sequence, using 12 pre-made trial sequences constructed by a Matlab program, using Gellermann (Gellermann, 1933) rules for an ‘appropriate’ randomization of psychophysical trials. The Gellermann rules are that the number of stimuli should remain the same to avoid stimulus bias, and that the stimulus is not presented more than three times in a row. The random schedules were adjusted so that there were never more than three trials in a row of either GO or NOGO trials.

Prior to each research session, the subject was voluntarily gated into the Pool C (Figure 1). A minimum of two researchers were present for each research session: The *experimenter* (K.A.H) and the *observer* (S.T.O., A.M. & A.H.). The *experimenter* conducted the trials and reinforced the seal following each correct response, while the *observer* recorded the progression of trials, quantity of reinforcement (number of fish pieces), the seal’s behavior during and between trials (i.e. response to a trial, swimming away from the station, or playing with fish), as well as any other relevant information concerning each trial or the session. The quantity of reinforcement remained constant throughout a single research session. The *experimenter* sat on the edge of the pool, above the test set-up, and the *observer* sat 1 meter away (Figure 2A). During each session, a quiet environment on or around the deck was made to minimize any distractions that may cause the animal to lose focus during a research session.

**Figure 2.**
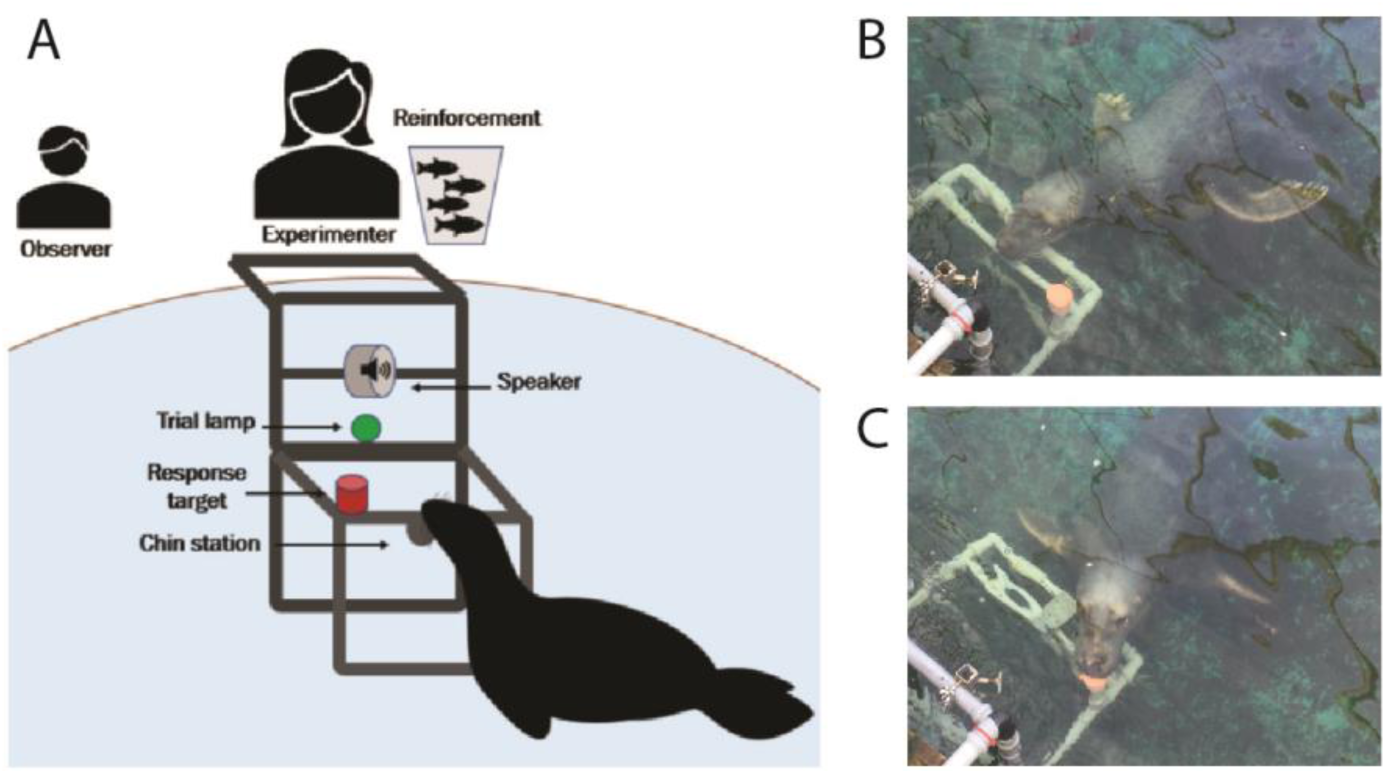
**A** Schematic of underwater setup for psychophysical research. Pictures depicting the positioning of the seal during the psychophysical testing. **B** Shows the animal positioned at the station, this is where the animal starts each trial, as well as indicates to trainer that it did not hear a tone, and **C** shows the animal signaling to the trainer that it heard a tone.

The station was located 75 cm below the surface and 50 cm in front of an underwater loudspeaker and a trial light (Figure 2A). Prior to the start of a trial, the subject was asked to place their chin onto a chin plate (Figure 2B). For a correct GO trial, the seal would leave the station upon the presentation of an audio signal and touch a red response target to the left of the station with its nose to indicate to the experimenter that it heard a tone (Figure 2C). For a correct NOGO trial, the seal would remain at the station until the end of the trial indicating that it did not hear any tone (Figure 2B). The trial began when the light turned on, signaling to the animal that an auditory signal may or may not be presented to them within the next 4 seconds. At the start of each session, the volume level of the auditory signal was always well above their hearing threshold. With each correct GO response, the signal intensity would decrease by 3dB, and with each incorrect GO response would increase by 3dB (Figure 3).

**Figure 3.**
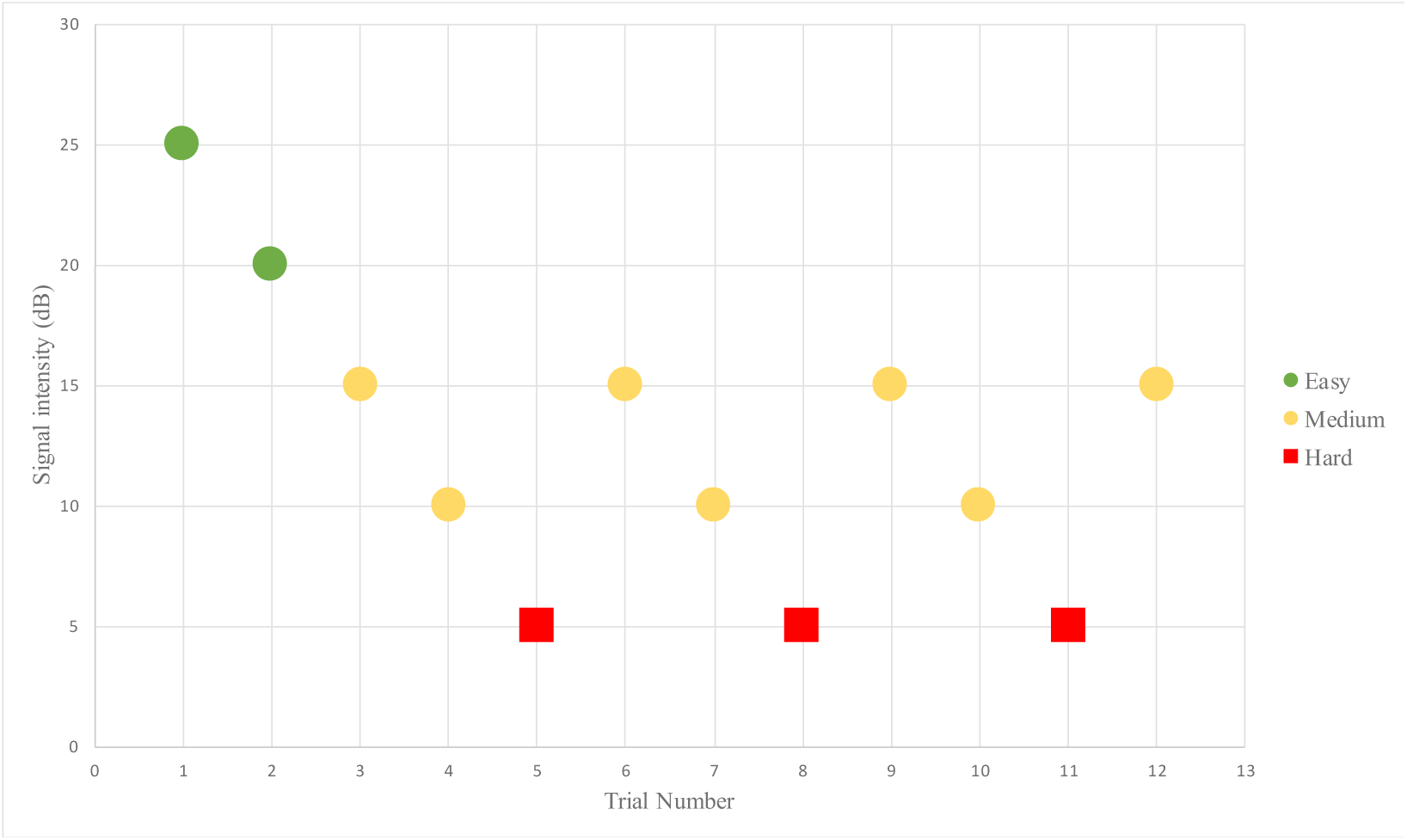
Example of one psychophysical research session. Green circles represent where the task is easy for the animal, and as the signal intensity decreases with correct go responses, the task becomes increasingly more difficult (red squares).

Each correct response was marked by an underwater buzz tone, a conditioned reinforcer, which indicated to the animal that it had responded correctly and to return to the trainer for reinforcement. The number and type of fish that the seal received varied between the individuals, but the quantity and type of fish remained constant between trials for an entire research session. For incorrect responses, the animal did not receive reinforcement and was asked to re-station for the start of the next trial. If the animal repeatedly left the station for a period of more than 10 seconds, a “time out” of one minute was given. Each research session had a predetermined number of trials to complete with no time limit to complete that task, but typically lasted 12-15 minutes. However, the session was ended if the inter-trial interval was too long and continued for consecutive trials (the animal swimming around the pool or playing with the fish reinforcement). Research sessions were conducted once per day, five times per week. These sessions were primarily conducted as the first session of the day, at 9am.

### Data analysis

A seal’s *level of motivation* towards the research session was determined by the frequency with which the animal left the station and swam away during a research session. Leaving station was defined as when an animal would leave the area around the chin station (>0.5 m) for longer than three seconds. In comparison, high motivation towards the research session was quantified as when the animal both stayed within a radius equal to or less than 0.5 meters from the chin station and returned to it within a time duration equal to or less than three seconds. The *interest in reinforcement* was determined by how long it took the seal to eat the reinforcement. If a seal took more than three seconds to eat the reinforcement or rejected the reinforcement (e.g. did not take the fish when it was presented, or dropped it after receiving it), it was interpreted as low interest in reinforcement. In addition to the observer’s notes, the daily caloric intake was calculated daily for each individual. It was then possible to calculate calories consumed during a research session, calories consumed as reinforcement as a percentage of the total caloric intake for the day, as well as a caloric value per reinforcement. Observations were taken from 83 sessions for M1, 86 sessions for M2 and 85 sessions for M3.

Data analysis was done in R version 2.4.0 (Team, 2017). We compared the behavior of the seals between the different subjects and over time (i.e. research sessions) using a General Linear Model (GLM) fitted with a Gaussian error structure. The response variable was seal behavior: (i) frequency of leaving the research station per session (which would indicate low motivation during the task) or (ii) playing with or rejecting fish (which would indicate low interest in reinforcement). The explanatory variables were the seal’s identity (M1/M2/M3) and the experimental session number (For M1 = 83 sessions, for M2 = 86 sessions and for M3 = 85 sessions).

We fitted a GLM with binomial error structure to test if the number of trials per subject was significantly different across a period of time (experimental sessions). The response variable was the number of trials accomplished per session and the explanatory variables were the seal’s identity and experimental session number.

## Results

All three seals left their chin stations during research sessions, but M1 left significantly more often than M2 and M3 (GLM testing, M1 vs. M2 SD = −15.32, d.f = 252, p < 2e-16; M1 vs. M3 SD = −13.71, d.f = 250, p = 3.81e-15; Figure 4). This means that the times that M2 and M3 stayed nearby the experimental set up was significantly higher than for M1 (Figure 4). M1 never needed more than three seconds to eat his reinforcement, never rejected it, and always showed a high interest in the reinforcement. For 100% of the sessions, M1 ate the reinforcement in less than three seconds, showing a high interest in his food reward. M2 showed lower interest in his food during research sessions and M3 showed a higher interest in reinforcement than M2 but significantly lower than M1 (GLM testing, M1 vs. M2 SD = 3.83, d.f = 252, p = 0.002; M1 vs. M3 SD = 2.52, d.f = 250, p = 0.043; Figure 5).

**Figure 4.**
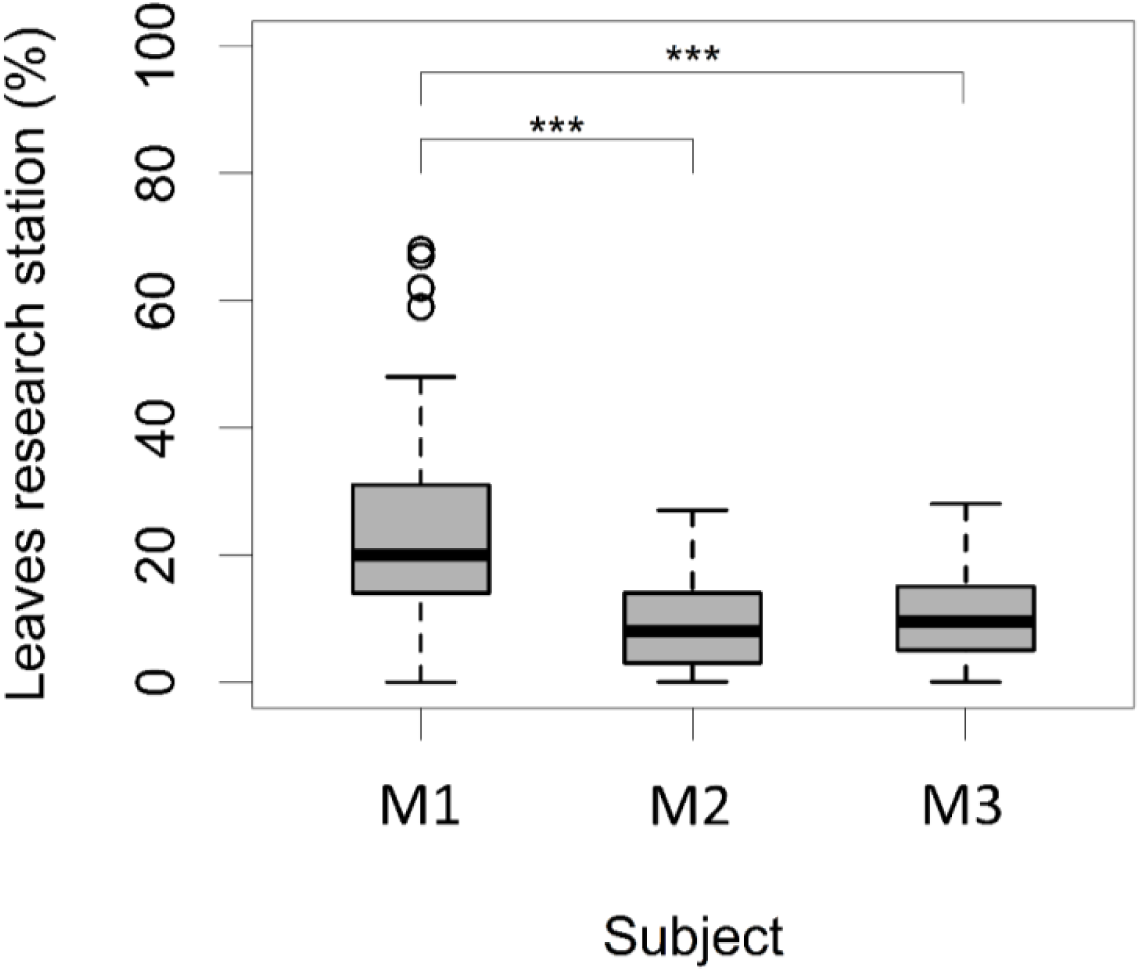
Number of times (%) each animal left the research station (greater than a 0.5 m radius) after each trial (N = 24 trials/session for M1; N = 34 trials/session for M2; N = 38 trials/session for M3, *** indicates p < 0.001, ** indicates p < 0.01, * indicates p < 0.05).

**Figure 5.**
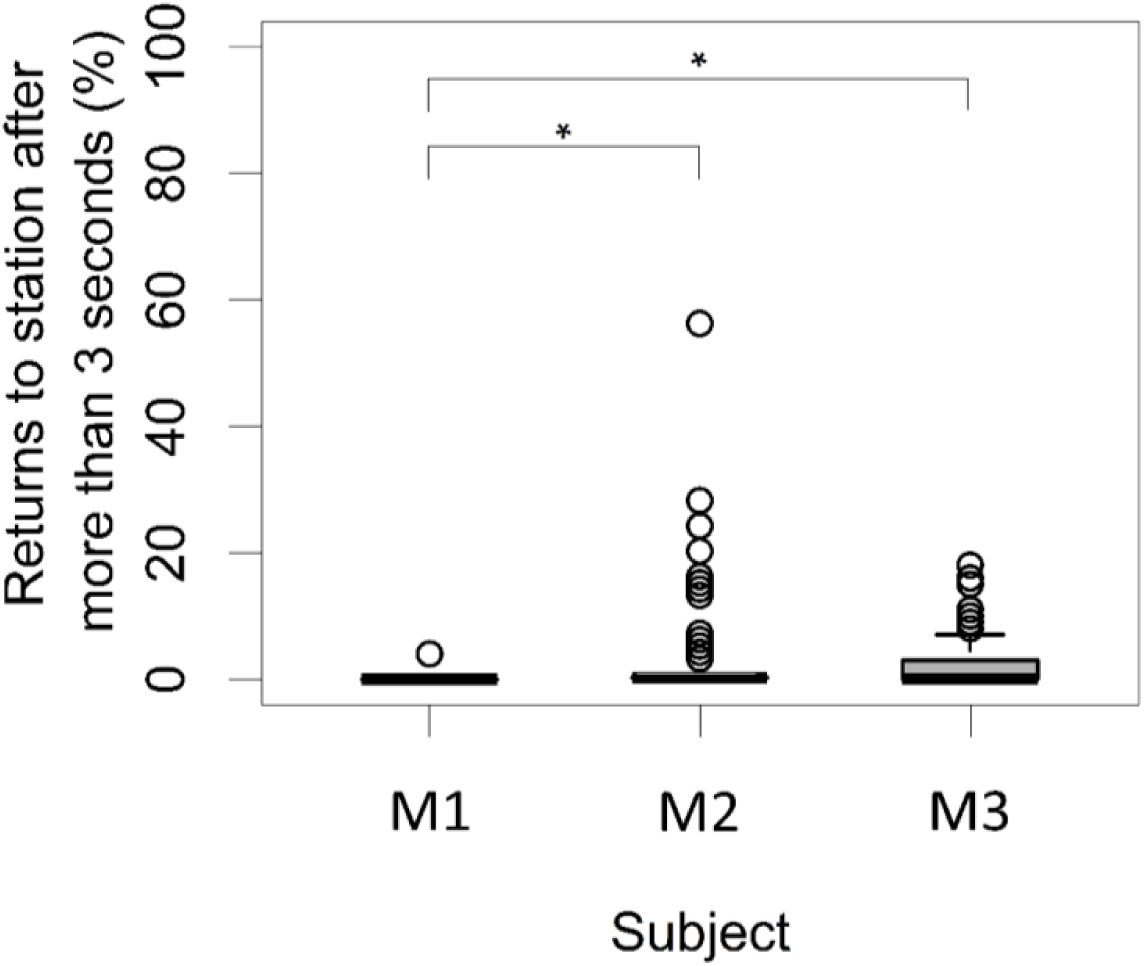
Number of times (%) each animal required more than three seconds to eat their reinforcement after each trial (N = 24 trials/session for M1; N = 34 trials/session for M2; N = 38 trials/session for M3, ** indicates p < 0.01, * indicates p < 0.05).

Regarding the number of trials accomplished per research session, there was no significant difference for either M1 (average 24 trials/session, GLM testing SD = −0.89, d.f = 82, p = 0.77) or M3 (average 38 trials/session; GLM testing SD = 0.76, d.f = 83, p = 0.32). For M2, there was a significant increase in the number of trials with time (average 34 trials/session; GLM testing SD = 3.17, d.f = 85, p = 4.6e-9). Therefore, there was a significant difference in the number of trials per session per animal (GLM testing SD = 0.69, d.f = 253, p = 2e-16) with the highest and most constant over time for M3, the lowest and least constant over time for M1, and from a low to a high number of trials for M2 over time.

There was a significant difference between the total amount of reinforcement received during a session between the three animals, which averaged 35% for M1, 22% for M2 and 46% for M3 of the individual’s daily caloric intake (GLM testing SD= 1.7, d.f = 252, p < 2e-11; Figure 6).

**Figure 6.**
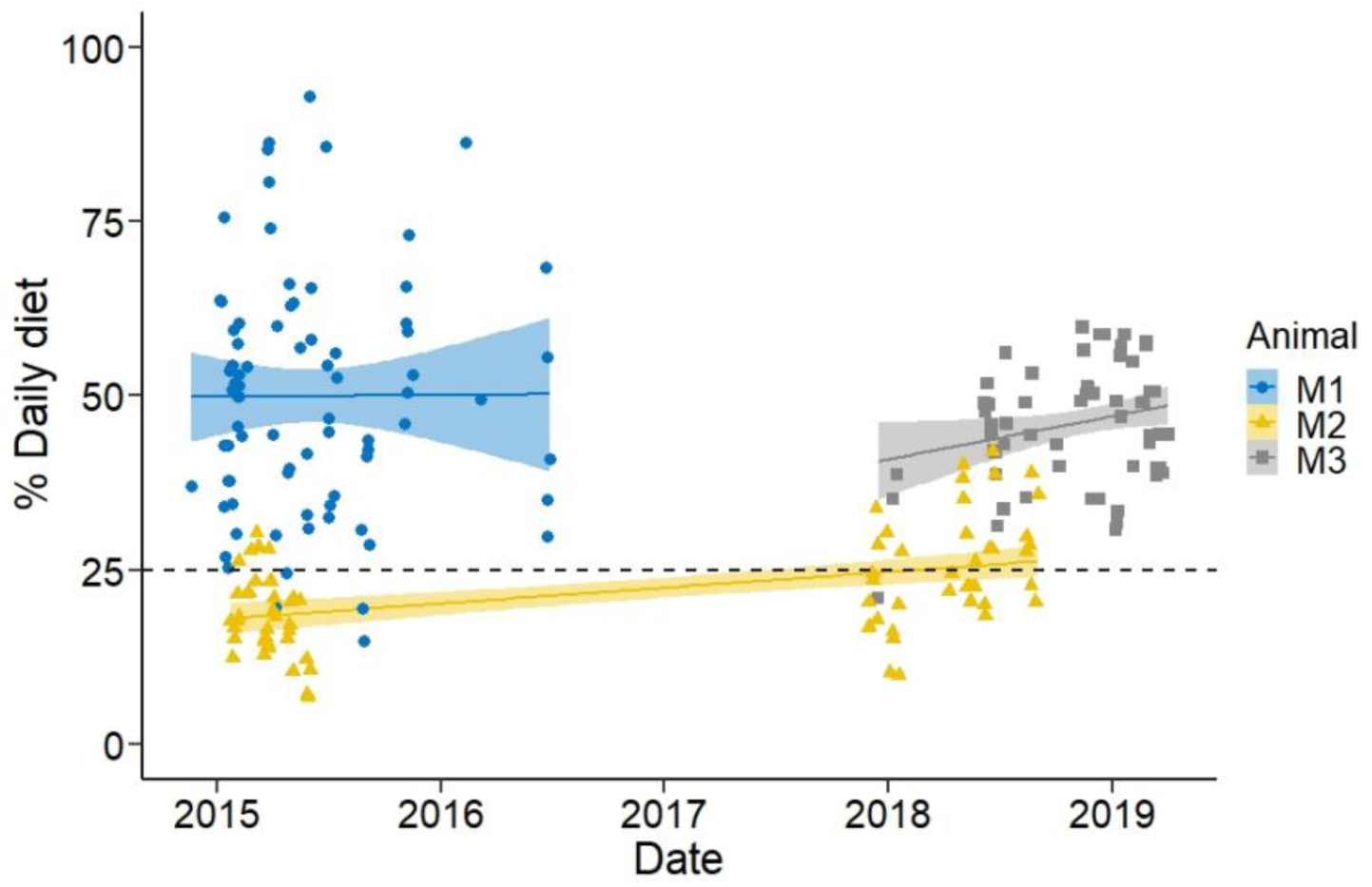
Percentage of the daily diet used as reinforcement per research session for each animal. Colored lines are derived from the linear model. The black dashed line is situated at the value 25%, as this represents the amount of food they would receive in a regular training session.

## Discussion

The goal of this study was to determine if the performance of three grey seals in psychophysical research could be used as a tool to provide positive enrichment. Psychophysical tasks have been known to provide cognitive challenges for animals (Keen et al., 2014; Stebbins, 2013). Our results demonstrate that participation in research can affect individuals in different ways.

Individual characteristics, such as personality and life experiences, can influence the animal’s performance when they face a cognitively challenging task. Motivation to participate in cognitive tasks have been shown in some great ape studies to be linked to different personality “traits”, such as audacity (Herrelko et al., 2012). Several variables have been identified to influence creative problem solving, such as openness, tolerance of ambiguity, divergent thinking, and intrinsic motivation (Davidson et al., 2003). Even though the seals studied participated in the research project, our results show that M1 was less motivated and required more reinforcement to participate. This may be due to differences in personality and life experiences between the three seals, whilst M2 and M3 were well-motivated participants in sensory response research. Such differences between the animals may have a large effect on their motivation to participate in research trials, as well as the enrichment they get out of such participation.

As an individual’s motivation to perform a research task may change the outcome of the study, the variation between the individual’s view on the significance of reinforcement may be attributed to the animal’s desire to participate in the trials. It is known that some animals are motivated to explore challenges without the need for extrinsic rewards (Clark et al., 2013). During the GO/NOGO testing, if the animal was to respond wrongly, no reinforcement was given. When the signal approaches the animal’s hearing threshold, the animal may not hear the stimulus even if it is present. From the viewpoint of the animal, it is not being reinforced despite having done well. This lack of reinforcement can cause stress or frustration. Considering the number of trials they participated in for each session, and the amount of reinforcement provided for each trial, we can conclude that M1 was showing more apathy or boredom to the task, and M2 and M3 were better at dealing with this frustration than M1, if this should be considered frustration by M2 and M3. In the case of M2, who would sporadically ignore reinforcement and return directly to the start position for the next trial. This suggests that this type of task is enriching for M2 (Langbein et al., 2009). In some of the cases, he would continue to participate in the session even though he had no interest in the reinforcement. This willingness to participate in research without reinforcement was never observed in M1, and rather required half of his total daily diet be used during these research sessions. In contrast to M1 and M2, M3 always showed interest in the reinforcement but also showed a high interest in the task. Previous research suggested that even within a single species, one type of enrichment often does not work for all individuals (Swaisgood & Shepherdson, 2005).

Their life histories and the environments in which these animals come from may provide additional information about the three seals and what may have influenced their approach to completing the research task. During M1’s first three years, he was not actively encouraged to be creative or introduced to new and challenging situations. In contrast, from a very young age, M2 and M3 were reinforced and encouraged for being creative and exhibiting play behavior, as well as being introduced to new and different situations.

The outcome of this study suggests that participation in research tasks can be considered enriching. It is beneficial to contemplate the personality of the animal involved (i.e. creativity, frustration tolerance, intrinsic motivation and curiosity) to ensure a positive outcome. Knowing that research can be used as an enrichment tool may increase the collaboration between research facilities, zoos and aquariums, provide a benefit to the animals under human care, and also contribute to our knowledge and understanding of wild species.

## Ethical Statement

The research followed all applicable international, national, and/or institutional guidelines for the care and use of animals. The animals were kept under Nature Protection Agency Permit SNS-342-00056 and Ministry of Food and Agriculture Permit 2300-50120-00003-09.

## Acknowledgements

The work is funded by grants from the Danish Council for Independent Research | Natural Sciences, the Carlsberg Foundation, and a grant from the Swedish Environmental Protection Agency. We would like to thank the many volunteers at the Marine Biological Research Centre for their assistance with data collection and observation during the underwater hearing sessions and Arturo Torres Ortiz for his help with statistics.

